# Human primary auditory cortex and insula encode perceptual decisions, not stimulus features

**DOI:** 10.64898/2026.07.03.736338

**Authors:** Camille Des Lauriers, Estelle Pruvost-Robieux, Marc Zanello, Cristina Filipescu, Alessandro Moiraghi, Elisabeth Landré, Johan Pallud, Gonzague Defrance, Tarek Sharshar, Jean-Julien Aucouturier, Martine Gavaret, Anaïs Llorens

## Abstract

The traditional view of perceptual decision-making assumes a largely feedforward cortical hierarchy, in which sensory regions encode stimulus features that are progressively integrated with top-down signals in higher order associative areas to guide decisions. However, recent work has cast doubt on whether stimulus-and decision-related signals actually follow such a predicted spatiotemporal organization, especially in naturalistic situations where sensory cues are subtle and decisions more strongly driven by internal strategies. Leveraging the unique spatiotemporal precision of human intracerebral recordings, we map here how stimulus and decision variables are represented along the auditory cortical hierarchy as patients engage in a social voice decision task with realistic, low-salience cues. Contrary to feedforward predictions, we found no clear spatial or temporal gradient separating stimulus- and decision-related effects; rather, decision effects emerged early during stimulus exposure and, strikingly, flowed back all the way to the most primary regions of the superior temporal gyrus. In addition, the direction of these early and primary decision signals closely reflected the variability in patients’ specific decision criteria. Taken together, these results strongly challenge the traditional feedforward model, supporting a view in which the activity of early auditory regions does not reflect subtle stimulus categories but is instead dynamically configured by task-dependent priors and response strategies.

**Significant Statement:** This work studies an ecological social-cognitive decision task based on subtle but natural vocal cues. Contrary to expectations, it shows that early and primary auditory activity in human intracerebral recordings does not reflect stimulus categories but instead patients’ decisions and decision criteria. These results are significant because they provide rare human intracerebral evidence that sensory regions are not static repositories of stimulus representations but rather dynamic, task-dependent filters that are configured by priors and decision criteria.

## Introduction

The traditional feedforward model of perceptual decision-making posits that sensory regions act as “bottom-up” feature extractors, passing increasingly abstract representations to higher-level associative areas responsible for computations like categorization and decision-making. In the auditory domain, this view dominates models of speech and auditory object categorization (Rauschecker and Tian, 2000; Hickok and Poeppel, 2004; Schirmer and Kotz, 2006), with processing streams extending anteriorly from primary auditory cortex (A1) toward lateral superior temporal, anterior temporal and prefrontal regions (Romanski et al., 1999).

However, growing evidence shows that sensory areas are not merely stimulus-driven but are also modulated by top-down signals, adjusting their tuning based on task demands (Gilbert and Li, 2013).

In primates, visual electrophysiology has shown that visual cortex (V1) receptive fields are modulated by spatial attention (Motter, 1993; Luck et al., 1997), and V2 neuronal gain is biased by perceptual expectations (Nienborg and Cumming, 2009). Predictive coding frameworks further propose that sensory regions may encode prediction errors rather than static stimulus representations (Friston, 2005), although direct electrophysiological evidence in humans remains limited (Solomon et al., 2021; Westerberg and Roelfsema, 2025).

Thus, sensory cortices may participate in both early feedforward analysis and in later feedback or decision-related processing (Cooper et al., 2023). How this balance unfolds in human sensory cortices during ecological perceptual decisions remains unclear. Several limitations have made this question difficult to address. First, most electrophysiological evidence for top-down influences comes from animal studies, with relatively little intracerebral human data. A recent intracerebral electroencephalography (iEEG) study in epileptic patients attempted to dissociate stimulus and decision features in human sensorimotor areas by contrasting report vs. no report tasks using identical tactile stimuli (Albertini et al., 2025). Report-specific electrodes were concentrated in associative regions ventral to primary sensorimotor cortex, where stimulus-specific electrodes were located, with later activity peaks, a pattern consistent with the hierarchical feedforward model. Whether a similar organization applies to processing streams with richer representational capacities, such as the auditory ventral stream, remains unknown. Second, many perceptual decision-making studies typically use suprathreshold psychophysical stimuli, which, while improving signal-to-noise ratios, conflate decision and stimulus features across trials (i.e., trials categorized as one percept also provide strong stimulus support for that category). They may also rely less on ingrained priors and response strategies than ecological, lower-salience social or emotional decisions (Shamay-Tsoory and Mendelsohn, 2019).

In this context, the superior temporal gyrus (STG) and the insula provide particularly informative regions of interest. The STG encompasses auditory cortical territories at the interface of dorsal and ventral processing streams and is centrally involved in the analysis of speech, voices, and socially relevant vocal cues (Rauschecker and Scott, 2009; Belin et al., 2011). The insula is well positioned to integrate sensory evidence with affective, interoceptive, salience, and decision-related signals (Menon and Uddin, 2010). Together, these regions allow us to test whether decision-related activity remains confined to higher order integrative cortices, or whether it also emerges within auditory sensory regions engaged by the task.

Here, we designed an ecologically relevant auditory decision task using low-salience vocal cues that were difficult to discriminate and required contextual or social inference. Unlike classical perceptual decision paradigms, in which highly salient stimuli produce strong stimulus-driven separability in sensory cortices, our task subtly embedded stimulus-category information, making perceptual reports more dependent on internal context and strategy. Leveraging the spatiotemporal precision of human intracerebral recordings, we used signal detection theory (SDT) to dissociate stimulus-category effects from response-category and decision criterion (*C*) effects across the auditory cortical hierarchy. Under a traditional feedforward model, stimulus-category effects should emerge early and dominate in auditory sensory regions, whereas choice-related effects should arise later and localize first to higher order associative areas. Contrary to this model, our results show that auditory sensory regions are not static repositories of stimulus representations, but dynamic, task-dependent filters that encode patients’ decisions and decision criteria more strongly than the underlying stimulus categories.

## Materials and Methods

### Patients

We recorded iEEG in N=23 neurosurgical patients with drug-resistant epilepsy (12 women; median age = 28 years old, range = 19-51 years old) who had been implanted with depth electrodes to define the epileptogenic network. Electrode implantation scheme was decided based on anatomo-electrical clinical correlations, with trajectory and anatomical coverage varying between patients according to presurgical needs. Both implantation and experimental procedures were carried out at the GHU Paris Psychiatry and Neuroscience Hospital. Of these 23 patients, 19 were retained for the present analyses because they had electrode coverage in the STG and/or the insula, the regions of interest in this study, as shown in Fig. 1B; detailed patient characteristics are reported in Table 1. The sample size is in line with standards in the field and comparable to, or larger than, that of previous intracerebral studies investigating intrinsic neural dynamics (Mercier et al., 2022; Cusinato et al., 2023).

**Figure 1:**
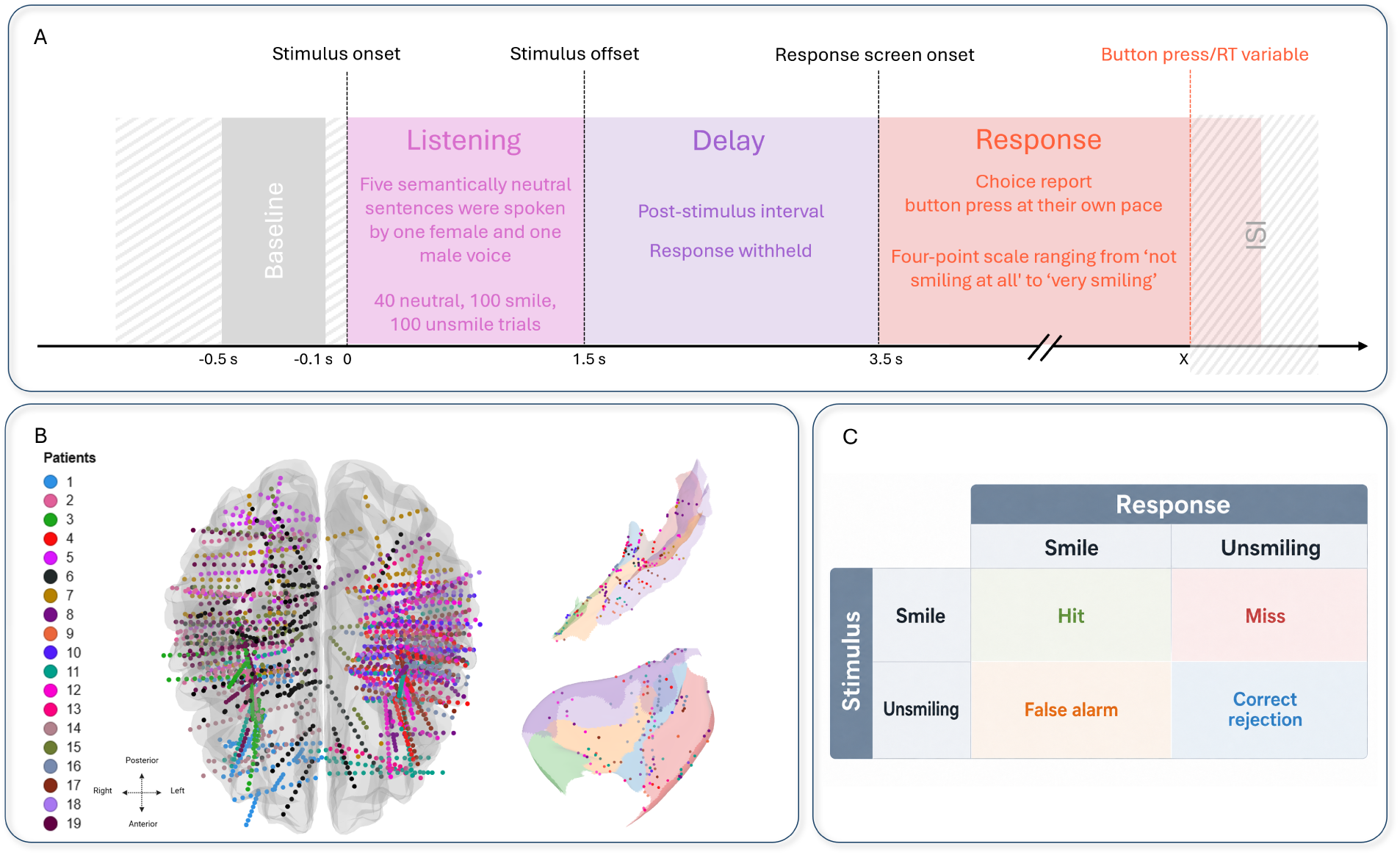
Experimental design, electrode coverage, and signal detection framework. **A,** Schematic representation of the experimental timeline. Trials included a prestimulus baseline period, followed by sentence presentation, a post-stimulus delay, and a response period. During listening, participants heard auditory sentences belonging to one of three stimulus categories: smile, unsmile, or neutral. After stimulus offset, participants withheld their response until the response screen appeared, then reported the perceived smiling content using a four-point scale ranging from “not smiling at all” to “very smiling”. **B,** Anatomical coverage of the retained iEEG contacts. Each dot represents one bipolar contact, colored according to patient identity. **C,** Signal detection theory was used to categorize behavioral responses: hits corresponded to smile stimuli reported as smiles, misses to smile stimuli reported as unsmiles, false alarms to unsmile stimuli reported as smiles, and correct rejections to unsmile stimuli reported as unsmiles.

**Table 1:** Patient sample and implantation coverage. Overview of the retained patient sample, with gender, age, iEEG implantation, and the number of implanted depth-electrode channels in STG and insula.

| Patient | Gender | Age (yo) | iEEG implantation | Number of depth-electrode channels |  |  |  |
| --- | --- | --- | --- | --- | --- | --- | --- |
|  |  |  |  | Left hemisphere |  | Right hemisphere |  |
|  |  |  |  | STG | Insula | STG | Insula |
| 1 | M | 19 | R insulo-fronto-orbital | 0 | 0 | 3 | 16 |
| 2 | F | 28 | R temporal | 0 | 0 | 34 | 21 |
| 3 | M | 20 | R fronto-parietal | 0 | 0 | 0 | 39 |
| 4 | F | 23 | L temporo-insular | 29 | 27 | 0 | 0 |
| 5 | F | 21 | R occipito-temporal | 0 | 0 | 8 | 0 |
| 6 | F | 51 | Bi-temporal | 7 | 2 | 16 | 1 |
| 7 | F | 30 | Bi-temporal | 15 | 0 | 19 | 1 |
| 8 | F | 23 | L temporo-insulo-parietal | 37 | 42 | 0 | 0 |
| 9 | F | 30 | L temporal | 20 | 6 | 0 | 0 |
| 10 | M | 25 | L temporal | 22 | 8 | 0 | 0 |
| 11 | F | 39 | Bi-temporal | 19 | 22 | 1 | 13 |
| 12 | M | 39 | L temporo-parieto-insular | 31 | 22 | 0 | 0 |
| 13 | F | 49 | L temporal | 34 | 11 | 0 | 0 |
| 14 | M | 26 | R insular | 0 | 0 | 15 | 29 |
| 15 | F | 21 | R fronto-temporal | 1 | 0 | 17 | 0 |
| 16 | F | 24 | L operculo-insular | 19 | 50 | 0 | 0 |
| 17 | M | 31 | L temporo-insular | 34 | 26 | 0 | 7 |
| 18 | M | 26 | L temporal | 27 | 8 | 0 | 0 |
| 19 | F | 43 | R fronto-temporo-parietal | 0 | 0 | 25 | 28 |

### Ethics statement

The study was approved by the French Comité de Protection des Personnes Sud-Est III (2020-A01340-39). All patients provided written informed consent prior to participation. The study is part of a clinical trial registered on ClinicalTrials.gov (NCT04810832).

### Stimuli

Auditory stimuli were generated from five neutral content sentences, adapted in the French language from (Russ et al., 2008). Each sentence was recorded by one woman and one man speaker, and then presented to patients in three versions: neutral, “smile” and “unsmile”. Smile and unsmile versions were created with a previously developed voice transformation algorithm, designed to simulate the ecological spectral changes that occur to human speech when a speaker is smiling (Arias et al., 2018). Specifically, the smile manipulation enhanced smile-related formant cues by shifting the first two formants upward and increasing energy in the upper-mid frequency region around the third formant, whereas the unsmile manipulation implemented the inverse transformation. These manipulations were validated as effective and natural in several previous studies (Arias et al., 2018; Arias et al., 2018), and notably leave the linguistic content and other vocal characteristics, such as pitch and speech rate, unchanged. All stimuli were edited to have exactly the same 1500 ms duration, and their loudness was normalized across all three stimulus categories (by equating the recordings’ root-mean-square intensity).

### Experimental design

The experimental timeline is summarized in Fig. 1A. Patients performed an auditory categorization task based on the smiling content of sentences. Each trial began with a 1500 ms listening period during which a sentence was presented, followed by a 2000 ms response-withholding delay, during which no response was allowed. This delay period was introduced to dissociate the electrophysiological recording of sensory processing from overt report. After 2000 ms, the response period started with the appearance of a four-point rating scale on the screen, ranging from “not smiling at all” to “very smiling” with two intermediate levels, with which patients were instructed to rate the smiling content of the recording. There was no time limit on the response duration. After the response, the next trial began following an interstimulus interval of 1750 ms, jittered at *±* 250 ms. The experimental session was organized into three blocks of 80 trials, separated by short pauses and was preceded by five practice trials to familiarize patients with the task. Across the whole session, 100 smile, 100 unsmile and 40 neutral stimuli were presented in a pseudorandom distribution such that the same sentence was not presented more than three times consecutively. Auditory stimuli were delivered through intra-auricular headphones and sound intensity was individually adjusted to a comfortable level for each patient.

### Behavioral data analysis

For analysis, response options 1-2 on the rating scale were grouped as the “unsmile” response category, and options 3-4 as “smile”. Based on this binarization, we then used SDT, illustrated in Fig. 1C, to categorize trials into hits (smile stimuli, judged as smile), misses (smile stimuli, judged as unsmile), correct rejections (CRs, unsmile stimuli, judged as unsmile), and false alarms (FAs, unsmile stimuli, judged as smile) (McNicol, 2005). Neutral trials were excluded from the analysis.

For each patient, we quantified decision criterion *C* as 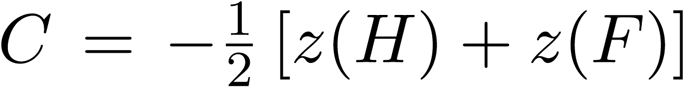, where *H* and *F* are the patient’s hit and false-alarm rates. Hit rate *H* was defined as the proportion of smile stimuli categorized as smile, 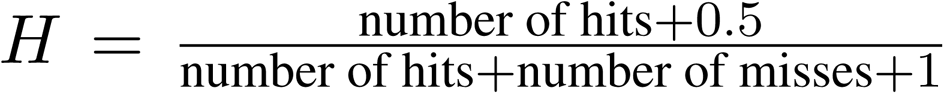, and FA rate as the proportion of unsmile stimuli categorized as smile, 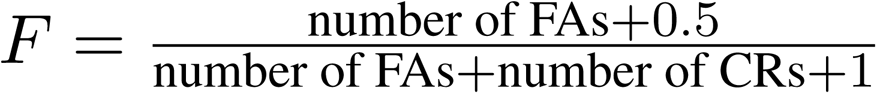. Both rates were corrected to avoid infinite *z* values when rates were equal to 0 or 1. By convention, negative values of *C* indicate a response bias toward smile reports, whereas positive values of *C* indicate a response bias toward unsmile reports. The decision criterion was therefore treated as a continuous behavioral measure indexing each patient’s position along the smile–unsmile decision criterion axis.

### iEEG recordings and preprocessing

iEEG recordings were acquired using a Micromed device (Natus, Pleasanton, CA, USA) from depth electrodes (ALCIS, Besançon, France or DIXI Medical, Besançon, France) implanted in various brain regions, with each electrode containing 5 to 18 platinum channels along its shaft. Electrodes were implanted using both orthogonal and oblique trajectories. When present, oblique trajectories provided denser sampling of the insular cortex than orthogonal trajectories. Data were acquired at 2048 Hz and downsampled offline to 1000 Hz. All channels were visually inspected prior to analysis and channels exhibiting persistent inter-ictal spiking, sustained epileptiform activity or excessive artifacts were excluded. Continuous recordings were then notch filtered at 50 Hz (and its subsequent harmonics) to remove line-noise contamination, and band-pass filtered between 0.1 and 250 Hz. iEEG signals were re-referenced in a bipolar montage to reduce the influence of a common reference and shared activity between neighboring channels, thereby favoring a more local representation of neural activity. This choice is classically motivated by the effects of volume conduction and common signals in intracerebral recordings, which are particularly relevant for the interpretation of focal analyses (Buzsáki et al., 2012). Moreover, this choice is relevant for high-frequency activity (HFA) analysis, as HFA is widely regarded as a relatively local marker of cortical processing (Lachaux et al., 2012). Peristimulus epochs were then extracted, spanning from −0.5 s before the stimulus onset to 4.8 s after its onset. All epochs were visually inspected, and any epochs with remaining artifacts were rejected. The baseline period of each epoch was defined as the interval [−0.5, −0.1] s preceding the stimulus. Processing of iEEG data was performed using MNE-Python (Gramfort et al., 2013).

### Electrodes localization and anatomical labeling

Post-implantation high-resolution head CT images were obtained and co-registered to pre-implantation MRI to determine the anatomical location of each iEEG channel (Filipescu et al., 2025). Image alignment and depth electrode leads localization were performed using the YAEL module of the RAVE (Reproducible Analysis and Visualization of iEEG) software (Wang et al., 2023). Cortical surface reconstruction and anatomical parcellation were performed within the same pipeline using the FreeSurfer software (Fischl, 2012), and channels were assigned anatomical labels according to the Destrieux cortical parcellation index (Destrieux et al., 2010). In this study, STG included both lateral superior temporal regions and transverse temporal/Heschl regions, allowing us to test whether decision-related effects extended into early auditory cortex.

Detailed anatomical coverage for each of the 19 patients is reported in Supplementary Table S1. Anatomical coverage varied across patients, as electrode implantation was determined exclusively by clinical requirements. Across retained patients, a total of 2304 channels located in grey matter were included in the whole-brain anatomical coverage analysis. At the region level, coverage was particularly dense in temporal and insular regions, which together accounted for 1711 of the 2304 channels, providing broad anatomical coverage for the regions investigated in the subsequent analyses. Within the regions of interest, 892 channels were assigned to the STG and/or the insula, comprising 487 channels in the STG and 405 channels in the insula. STG coverage was available in 18 patients, insular coverage was available in 17 patients, and 16 patients had coverage in both regions. Coverage was distributed across both hemispheres, with 13 patients contributing left-hemisphere STG channels, 9 contributing right-hemisphere STG channels, 11 contributing left-hemisphere insular channels, and 9 contributing right-hemisphere insular channels.

### HFA estimation

HFA was estimated from bipolar iEEG signals using Morlet wavelet decomposition of single trials over the 60–150 Hz range, with 28 logarithmically spaced frequencies and a number of cycles increasing with frequency from 5 to 12. To reduce boundary effects related to the wavelet transform, a guard interval was applied at both edges of the epoch, scaled to wavelet duration, and only the retained time samples entered subsequent analyses. Power estimates were computed on an extended time window (−1.5 to 5.8 s) to limit edge artifacts. HFA was then log-transformed and baseline-centered at the single-trial level by subtracting the mean log-power over the prestimulus baseline period (−0.5 to −0.1 s). Trials were then cropped to the analysis window (−0.5 to 4.9 s). HFA was obtained by averaging power across frequencies within the 60–150 Hz band, excluding narrow ranges around line-noise harmonics (98–102 Hz and 148–150 Hz).

#### Identification of task-responsive channels

We first identified channels that were task-responsive, independently of the experimental contrasts of interest and regardless of the condition. Task-responsive channels were defined as channels showing significant task-related modulation of HFA relative to baseline. Channels showing a significant effect after correction were classified as task-responsive, and retained for subsequent analyses.

#### Effects of stimulus-category and response-category

All analyses related to effects of stimulus-category and response-category were restricted to bipolar channels that both had at least one channel in gray matter and were identified as task-responsive by the procedure above.

Several types of contrasts were examined. First, stimulus-category and response-category effects were assessed using overall category contrasts. Stimulus-category effects were assessed by comparing trials with smile and unsmile stimuli, irrespective of the participant’s behavioral response. Response-category effects were assessed by comparing trials reported as smile with trials reported as unsmile, irrespective of the stimulus category. These overall contrasts therefore provided a broad assessment of stimulus-related and response-related activity across the task. To further dissociate stimulus-category and response-category effects, neural activity was also compared across SDT categories. Contrasts between hits and FAs, and between misses and CRs, were interpreted as stimulus-category effects with response category held constant.

Conversely, contrasts between hits and misses, and between FAs and CRs, were interpreted as response-category effects with stimulus category held constant (see e.g. (Arias et al., 2018) for a similar analysis strategy). Thus, the SDT analysis was used to test conditional stimulus-category and response-category effects that may not be apparent in the overall category contrasts.

Because response-category effects may reflect not only the reported category but also each patient’s individual response strategy, we also examined whether the direction of significant response-related HFA effects was aligned with, or opposite to, the patient’s decision criterion *C*. The subject-level classification and statistical testing procedure used for this analysis are described in the Statistical analyses section.

### Statistical analyses

Unless otherwise specified, all neural analyses were two-sided and were performed on baseline-corrected HFA time courses. Statistical testing was restricted to the three task periods defined by the experimental design: the listening period, corresponding to stimulus presentation (0-1.5 s after stimulus onset); the response-withholding delay (1.5-3.5 s); and the response period, starting at response-scale onset (3.5-4.8 s after stimulus onset).

#### Task responsive channels

Task responsiveness was assessed for each channel and each task window using a one-sample cluster-based permutation test against zero across trials (Maris and Oostenveld, 2007). The cluster-forming threshold was set to *p <* 0.01 and converted to the corresponding t-threshold according to the number of trials available for the tested subject. Cluster significance was evaluated using 5000 permutations, and only clusters reaching cluster-level *p <* 0.01 and lasting at least 50 ms were retained. For each channel and each task window, we retained the p-value of the most significant valid cluster, defined as the valid cluster with the lowest cluster-level p-value. If no valid cluster was found, the corresponding lead × window test was assigned a p-value of 1. These channel by window p-values were then submitted to false discovery rate (FDR) correction with *q <* 0.05, applied jointly across all channel by window tests within each subject. Channels showing a significant effect in at least one window after correction were classified as task-responsive.

#### Effects of stimulus-category and response-category

Condition-related effects were assessed separately for each tested channel and each task window using two-sample cluster-based permutation tests across time. This same statistical framework was applied to all condition-related contrasts, including stimulus-category contrasts, response-category contrasts, and SDT category contrasts. The cluster-forming threshold was set to *p <* 0.05 and converted to the corresponding t-threshold according to the number of trials available in the two conditions being compared. Cluster significance was evaluated using 5000 permutations, and only clusters reaching cluster-level *p <* 0.05 and lasting at least 25 ms were retained. Analyses were performed only when at least 8 trials per condition were available. For each channel and each window, the minimum valid cluster *p* value was retained. These values were then submitted to FDR correction with *q <* 0.05, applied separately within each task window for each contrast.

#### Analysis of response-related effects relative to decision criterion

To analyse how response-category effects on HFA activity relate to individual behavioral strategies, we restricted the analysis to patients who had at least one electrode channel significant for a response-category effect (i.e. in either the hit vs. miss or the FA vs CR contrast) in any of the three task windows. These patients were categorized as having smile-directed or unsmile-directed activity, using the following procedure: first, each significant channel of the patient was labeled as smile- or unsmile-directed, according to whether HFA activity in that channel was larger in hit than miss and/or larger in FA than CR (smile-directed) or the opposite (unsmile-directed); second, the patient was categorized as smile- or unsmile-directed according to the majority label of their channels. That analysis was done separately using electrode channels in the STG and the insula regions.

We then visualized and analyzed whether the direction of the response-category effect reflected individual-level decision criteria *C*. We described patients as “aligned with criteria/bias” when higher HFA was observed for the response category favored by the patient’s *C*, namely smile-directed HFA in patients favoring the smile-response category (i.e. *C <* 0), and unsmile-directed in patients with *C >* 0. We tested for a systematic association of HFA direction and criteria using Fisher exact tests on the 2 *×* 2 contingency table assigning patients to either of the criteria and HFA directions. HFA ties, i.e. patients with equal numbers of smile-directed and unsmile-directed electrode channels, were removed from the analysis.

## Results

### The auditory decision task elicited individually variable response strategies

Patients performed the auditory categorization task by rating the smiling content of each sentence on a four-point scale. Responses were subsequently binarized into smile and unsmile reports, allowing trials to be classified according to SDT as hits, misses, CRs and FAs.

Across the 19 retained patients, a total of 3364 trials were available after exclusion of neutral trials and artifact-contaminated trials, including 1671 smile trials and 1693 unsmile trials. Importantly, the task yielded usable trial counts in all four SDT categories, with a median of 56 hits per patient [range: 14–79], 37.5 misses [range: 13–79], 49 correct rejections [range: 12–81], and 42.5 false alarms [range: 18–82]. This distribution indicated that both correct and incorrect categorizations were sufficiently represented, allowing subsequent neural analyses to compare HFA activity across SDT categories.

Across all retained patients, task performance was numerically above chance but did not reach the conventional significance threshold (accuracy: M = 55.67%, SD = 12.09; chance level: 50%; *t*(18) = 2.04, *p* = .056; *d′*: M = 0.28, SD = 0.68; *t*(18) = 1.77, *p* = .093), consistent with the intended ambiguity of the task. In the subset of patients contributing to the decision-related electrode and bias analyses, performance was significantly above chance (accuracy: M = 57.92%, SD = 7.77; *t*(8) = 3.06, *p* = .016; *d′*: M = 0.42, SD = 0.41; *t*(8) = 3.05, *p* = .016), indicating that these analyses were based on patients who showed measurable, although modest, task sensitivity.

Signal detection analysis showed that patients varied in their decision criterion. Median decision criterion *C* was −0.02 [range: −0.51–1.51]. By convention, negative values of *C* indicate a response tendency toward smile reports, whereas positive values indicate a response tendency toward unsmile reports. Accordingly, 13 patients showed a response tendency toward smile reports, and 6 patients showed a response tendency toward unsmile reports.

Thus, despite performing the same auditory decision task, patients differed in their position along the smile–unsmile decision axis. This inter-individual variability in decision criterion was used in subsequent analyses to interpret the direction of neural effects relative to each patient’s response tendency. Detailed individual behavioral measures are reported in Supplementary Table S2.

### Task-responsive HFA activity is broadly distributed, prominent in the STG and insula, and predominates during listening

We first identified task-responsive channels independently of the experimental contrasts of interest, as shown in Fig. 2, by testing for significant HFA modulation relative to the prestimulus baseline during the listening, delay, or response periods.

**Figure 2:**
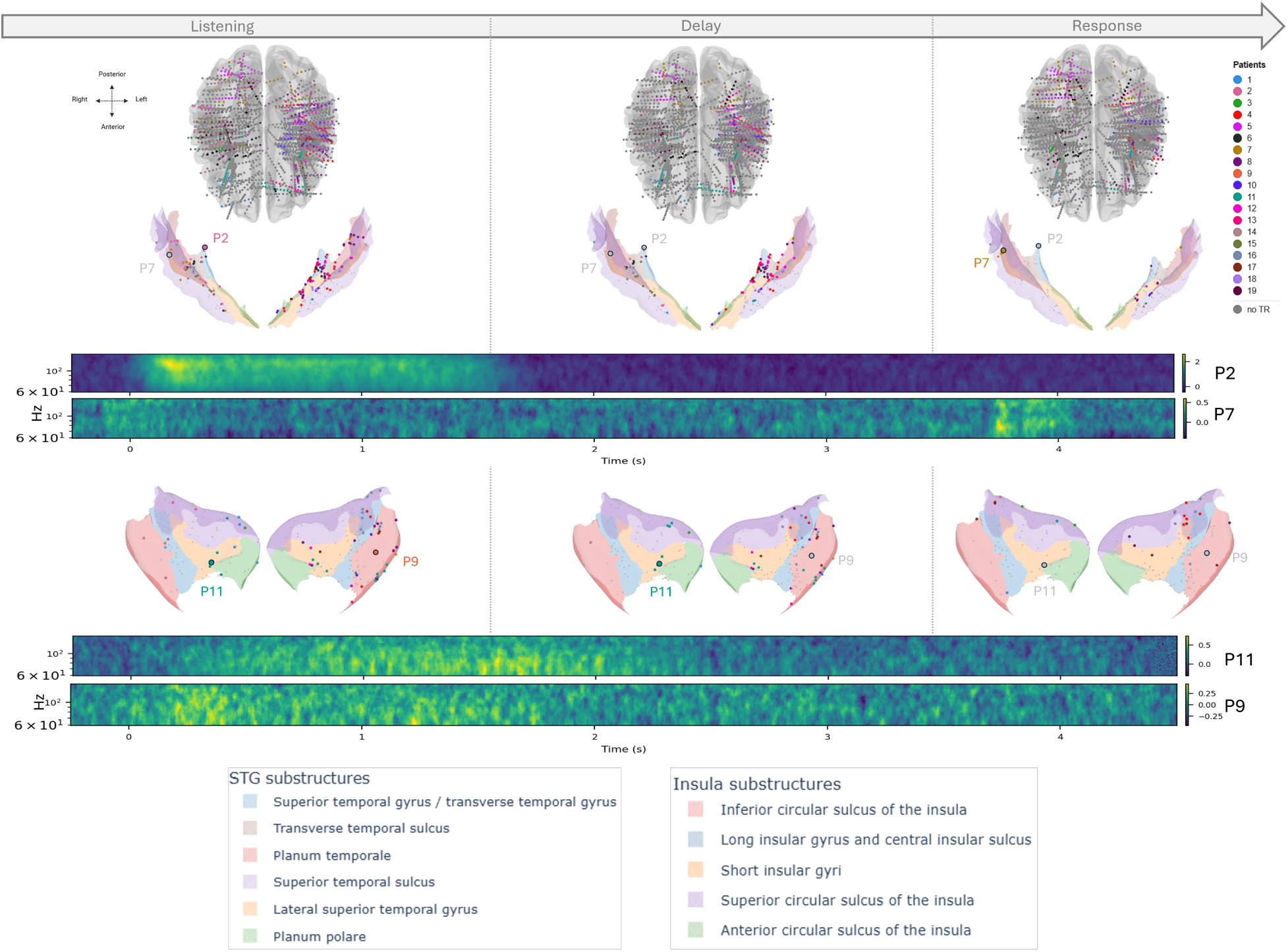
Task-responsive HFA activity in STG and insula. **Top:** Whole-brain top views provide an overview of the anatomical distribution of implanted bipolar contacts across patients during each task period. **Middle and bottom:** Task-responsive bipolar contacts are then shown for the STG (middle panels) and the insula (bottom panels) during listening, delay, and response periods (left to right). Colored contacts indicate task-responsive channels and are color-coded by patient identity; grey contacts indicate non-task-responsive channels. Cortical projections show the corresponding STG and insular substructures. Representative time-frequency maps from selected contacts illustrate task-related HFA modulation across the task and correspond to example electrodes from patients P2 and P7 for the STG (middle), and P11 and P9 for the insula (bottom).

This analysis revealed that the task engaged a broad network of brain areas. Across the retained grey-matter channels, 1016 channels showed significant task-related HFA modulation in at least one task window, corresponding to 44.1% of tested channels. Task-responsive channels (Fig. 2-top) were distributed across temporal, insular, frontal, parietal, and occipital regions, but were particularly concentrated in temporal and insular cortices, which together accounted for 71% of all task-responsive channels. Task responsiveness was slightly higher in the left hemisphere, involving 45.9% of tested channels compared with 43.4% of tested right-hemisphere channels.

We then focused on the regions of interest. In the STG (Fig. 2-middle), 296 out of 487 tested channels were task-responsive, corresponding to 60.8%. Task responsiveness in the STG was broadly bilateral, with comparable proportions of task-responsive channels in the left and right hemispheres. This bilateral distribution is important because the present task involved both vocal and socially meaningful information, and therefore cannot be reduced to a simple left-hemisphere speech account or to a right-hemisphere emotion/prosody account (Schirmer and Kotz, 2006). In the insula (Fig. 2-bottom), 158 out of 405 tested channels were task-responsive, corresponding to 39.0%, with a higher proportion of task-responsive channels in the left than in the right hemisphere. Detailed proportions by region, hemisphere, and task period are reported in Table 2.

**Table 2:** Pooled proportion of task-responsive channels in the STG and insula by hemisphere and task period. TR indicates task-responsive channels; TR any refers to channels showing significant task-related modulation in at least one task period. Values are reported as percentages with raw counts in parentheses.

| ROI | Hemisphere | Subjects | Tested channels | TR any | TR listening | TR delay | TR response |
| --- | --- | --- | --- | --- | --- | --- | --- |
| STG | Left | 13 | 325 | 59.4% (193/325) | 56.3% (183/325) | 42.8% (139/325) | 7.4% (24/325) |
| STG | Right | 9 | 162 | 63.6% (103/162) | 59.9% (97/162) | 32.1% (52/162) | 6.8% (11/162) |
| STG | Bilateral | 18 | 487 | 60.8% (296/487) | 57.5% (280/487) | 39.2% (191/487) | 7.2% (35/487) |
| Insula | Left | 11 | 238 | 42.4% (101/238) | 33.2% (79/238) | 24.8% (59/238) | 13.0% (31/238) |
| Insula | Right | 9 | 167 | 34.1% (57/167) | 26.3% (44/167) | 10.2% (17/167) | 7.2% (12/167) |
| Insula | Bilateral | 17 | 405 | 39.0% (158/405) | 30.4% (123/405) | 18.8% (76/405) | 10.6% (43/405) |

Across time, task responsiveness in STG and insula was distributed across the three task periods, with a larger occurrence during listening. Across both regions, 381 channels were task-responsive during listening (0-1.5 s), 251 during the delay period (1.5-3.5 s), and 78 during the response period (3.5-4.8 s). These window-specific counts were not mutually exclusive, because a given channel could be task-responsive in more than one period.

These task-responsive channels corresponded to a set of 18 patients in the STG, and 16 patients in the insula, which were kept for subsequent analysis. The proportion of task-responsive channels by patient, hemisphere, and task window is reported separately for the STG and insula in Supplementary Table S3. This independently defined task-responsive set was used as the anatomical and functional basis for all subsequent condition-related analyses.

### STG and insula show stronger response-category effect than stimulus-category effects

We next tested whether task-responsive HFA activity differentiated stimulus-category and response-category using overall contrasts. Stimulus-category effects were assessed by comparing smile and unsmile stimuli, irrespective of the patient’s response, whereas response-category effects were assessed by comparing trials reported as smile versus trials reported as unsmile, irrespective of the stimulus category.

When tested using this overall stimulus-category contrast, stimulus-category effects were limited. Across the STG and insula, significant smile-versus-unsmile stimulus effects were observed in a mere 10 channels from 5 patients (STG: 8 channels/3 patients; Insula: 2 channels/2 patients).

In contrast, response-category effects were more frequently observed. Across the regions of interest, significant smile-report-versus-unsmile-report effects were observed in 70 channels from 15 patients (STG: 57/11; Insula: 20/10).

Thus, in these overall category contrasts, condition-related HFA activity in the STG and insula was more strongly associated with the patient’s response than with the stimulus category alone. However, because these overall contrasts ignore the other task dimension, they may miss effects that depend on the joint configuration of stimulus category and response – a possibility that must be considered because of the large variety of decision criteria across participants. This pattern motivated a signal-detection-based analysis aimed at dissociating stimulus category from behavioral report more explicitly.

### Response-related effects emerge early and in primary sensory areas, contradicting the hierarchical information-processing view

Because the overall category contrasts collapsed across the other task dimension, we next used the SDT decomposition to test conditional stimulus- and response-category effects. In Fig. 3, we visualize these effects in the form of a square, where vertical edges correspond to stimulus-category effects (hit vs FA; miss vs CR) and horizontal edges correspond to response-category effects (hit vs miss; FA vs CR).

**Figure 3:**
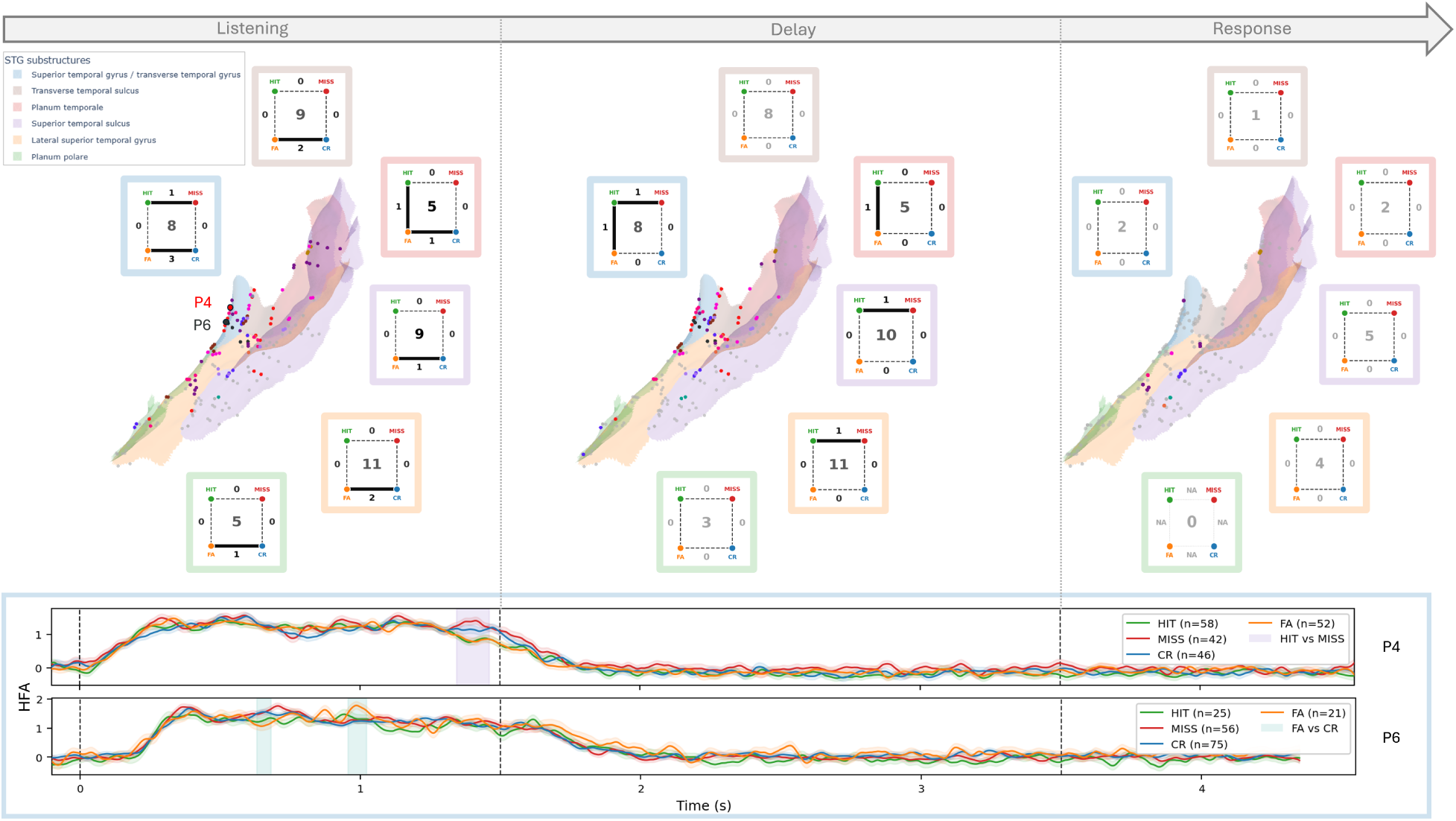
Response-related effects on HFA activity emerge early and upstream across the subregions of the superior temporal gyrus. **Top:** Significant contrasts between SDT response categories, grouped across participants for 6 STG subregions and the 3 temporal periods of the task. For each subregion and period, effects are visualized in the form of a square, where vertical edges correspond to stimulus-category effects (hit vs FA; miss vs CR) and horizontal edges correspond to response-category effects (Hit vs miss; FA vs CR). Numbers on the edges indicate the number of patients with at least one significant channel for the corresponding SDT contrast. The central number inside the SDT square indicates the number of patients with at least one task-responsive channel in that subregion and period. SDT effects were dominated by response-related horizontal edges and, contrary to what could be predicted by the classical hierarchical information-processing view, were mainly observed early (listening period) and upstream all the way to primary auditory regions such as the superior temporal/transverse temporal gyrus and the transverse temporal sulcus. **Bottom:** HFA time courses for two representative channels, indicated by larger circles in the top panel. Curves represent single-participant activity averaged over hit, miss, FA, and CR trials, and confidence intervals indicate the variability of activity across trials, within-participant. Vertical colored bands indicate time intervals showing significant differences, according to cluster-based permutation tests.

This analysis confirmed that HFA activity in the STG and insula was predominantly associated with the patient’s response rather than with the stimulus category. Across the regions of interest, significant SDT-edge effects were observed in 48 channels from 10 patients (STG:34/7; insula: 17/8). Most of these effects involved horizontal edges of the SDT square, corresponding to response-category differences with stimulus category held constant. Horizontal response-related effects were observed in 39 channels from 9 patients (STG: 30/6; insula:11/8), whereas vertical stimulus-related effects were only observed in 13 channels from 6 patients (STG: 7/4; Insula: 7/3).

Importantly, response-related effects were not associated with later time periods, nor were they secondary to prior stimulus-related effects, as would be predicted by a classical hierarchical information-processing view. First, there was very little overlap between both types of effects, with only 4 channels showing both types of effects across our dataset. Second, the majority of significant horizontal-edge effects were observed during the listening period (31 channels /7 patients), and their occurrence decreased during the delay period (9 channels / 5 patients) and was almost absent in the response period (1 channel / 1 patient). This temporal profile notably indicates that response-related HFA activity emerged before overt report execution and was therefore unlikely to reflect only late motor or response-production processes.

Another prediction of the classical hierarchical information-processing model is that stimulus-related effects would occur upstream in primary sensory areas, and decision-related effects would occur downstream in areas linked to associative processing. Strikingly here, not only were stimulus effects largely absent from the primary auditory cortex, but decision effects flowed upstream all the way to primary sensory regions. In the STG (Fig. 3), decision effects occurred notably in the superior temporal/transverse temporal gyrus and the transverse temporal sulcus (corresponding to Heschl’s region and adjacent early auditory cortex), and showed no particular prevalence in associative regions such as the lateral superior temporal gyrus or planum temporale.

A similar pattern was observed in the insula. While not generally considered a primary auditory area (but see (Berger et al., 2026)), the insula is generally considered to obey a posterio-anterior axis of information processing, with posterior insula showing auditory responses that resemble those observed in Heschl’s gyrus (Zhang et al., 2019; Berger et al., 2026) (see also Fig. 2-bottom), and anterior areas being associated with decision-making (Loued-Khenissi et al., 2020; Llorens et al., 2023). Here again though, our data strongly contradicted this common view: as shown in Fig. 4, decision effects occurred early and flowed back all the way to posterior regions in the inferior circular sulcus, long insular gyrus and central insular sulcus.

**Figure 4:**
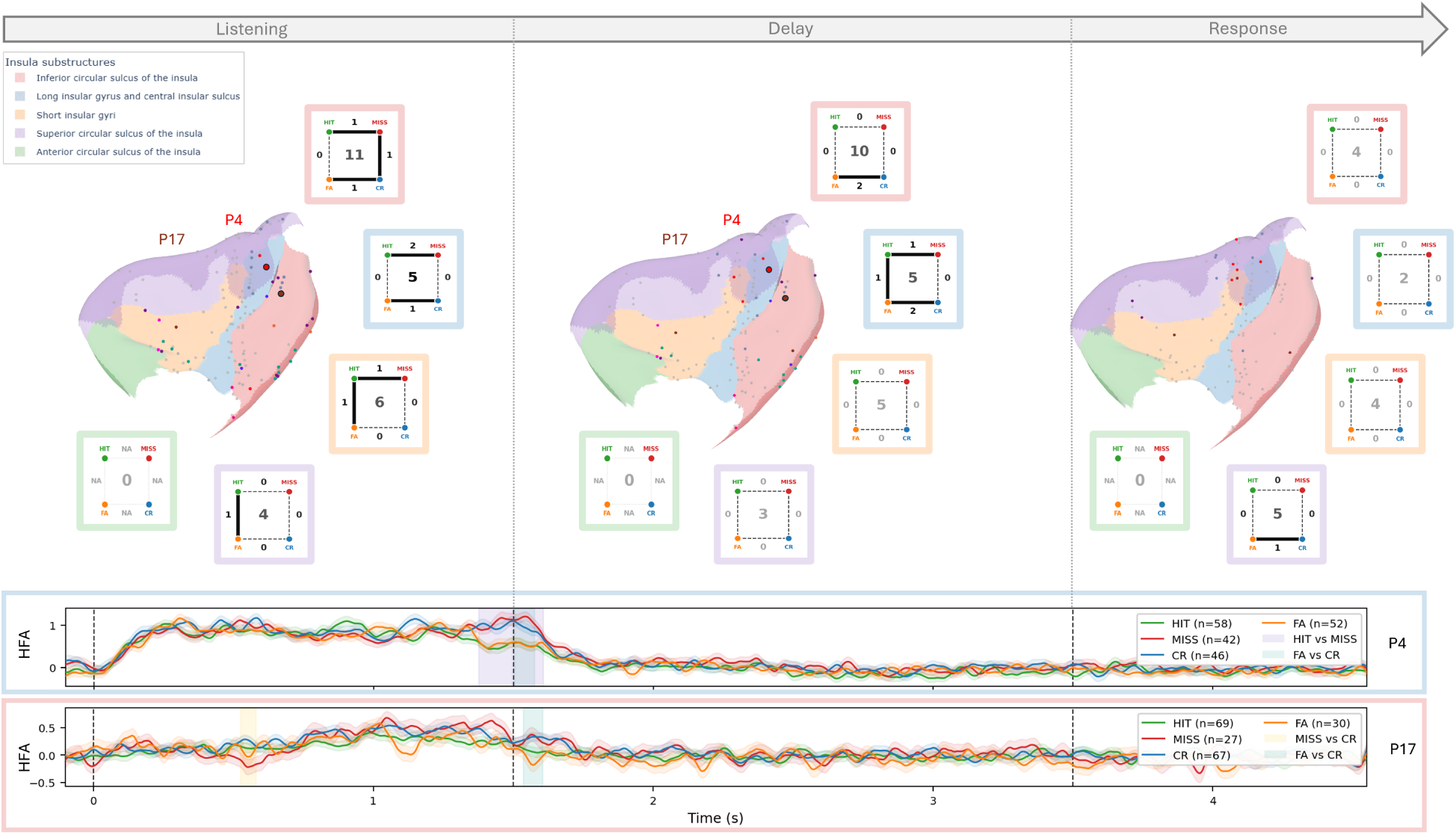
Response-related effects on HFA activity in the insula occur early and flow back all the way to posterior subregions. **Top:** Significant contrasts between SDT response categories, grouped across participants for 5 insular subregions and the 3 temporal periods of the task. For each subregion and period, effects are visualized in the form of a square, with the same organization as Fig. 3. As in the STG, SDT effects were dominated by decision contrasts (horizontal edges), occurred early and flowed back all the way to posterior regions in the inferior circular sulcus, long insular gyrus and central insular sulcus. **Bottom:** HFA time courses for two representative channels, indicated by larger circles in the top panel. Curves represent single-participant activity averaged over hit, miss, FA, and CR trials, and confidence intervals indicate the variability of activity across trials, within-participant. Vertical colored bands indicate time intervals showing significant differences, according to cluster-based permutation tests.

### The direction of decision effects reflects individual decision criteria

Finally, we asked whether the direction of decision effects (i.e. whether hit *>* miss, FA *>* CR, or the opposite) has any relation to individual decision criteria, as measured with SDT analysis of participant behaviour (see *Methods*).

In both STG and insula, there was a consistent negative association between the direction of response-related effects and individual decision criteria, in which higher HFA was seen for response categories that *opposed* the participant’s bias (e.g. hit*>*miss and FA*>*CR HFA activity in participants who favor the “unsmile’ response option).

As shown in Fig. 5-top, the direction of response-related effects in the bilateral STG was opposite to decision criteria in 4 out of 6 patients (left STG: 4 to 1, right STG: 1 to 1) – an association which, given the small sample size, did not reach statistical significance (Fisher exact test: *p* = 0.467). Similarly, as shown in Fig. 5-bottom, response-related effects in the bilateral insula were in the opposite direction to decision criteria for 6/7 patients (left: 4 to 1, right: 2 to 0) – an association which, again, did not reach statistical significance (Fisher exact test: *p* = 0.143).

**Figure 5:**
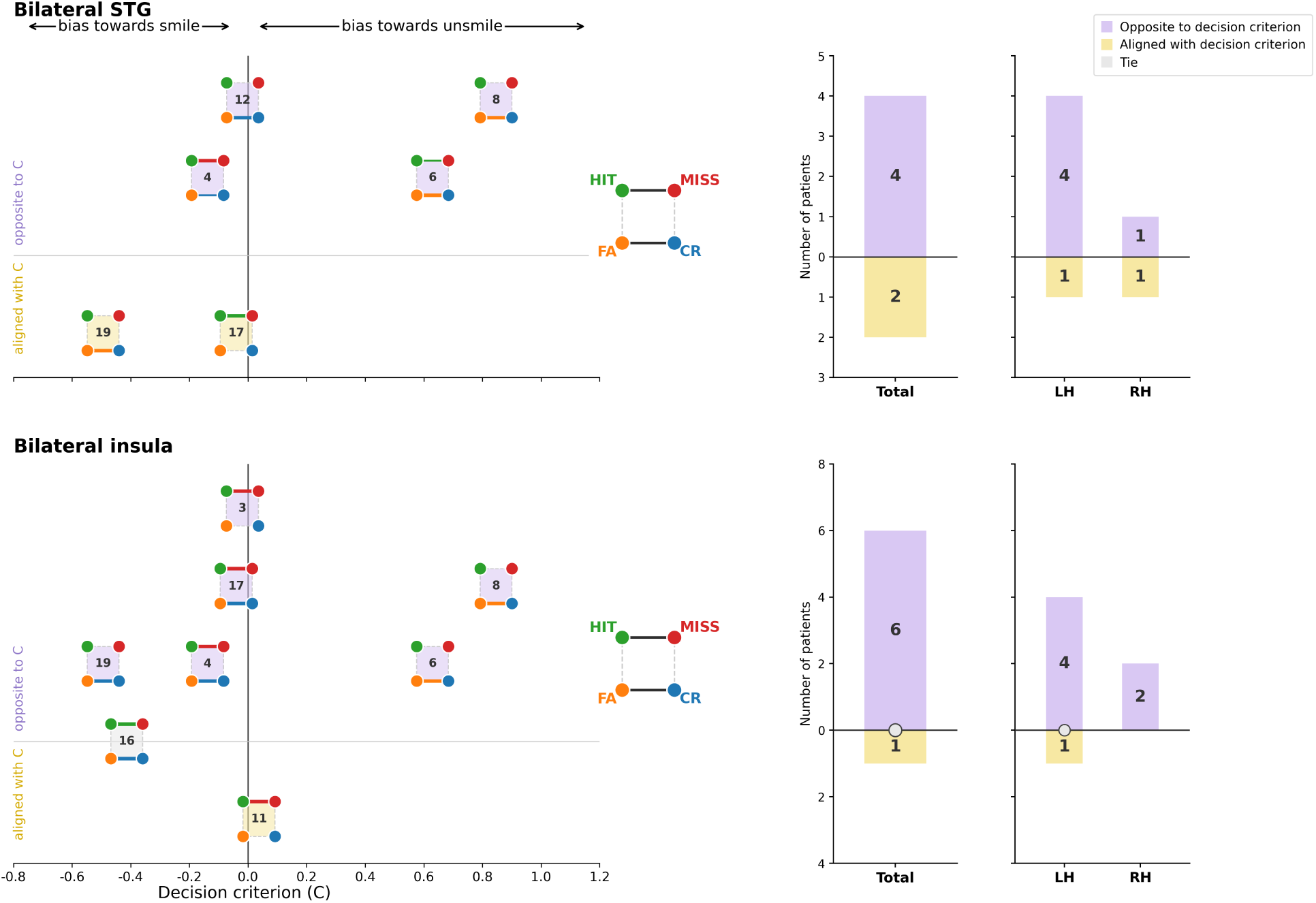
A consistent negative association between individual decision criteria and the direction of response-related effects on STG and insula HFA. Results are shown separately for the bilateral STG (**Top**) and the insula (**Bottom**). In each region, patients with significant decision effects are represented as individual SDT squares (the number inside the square indicates the patient identifier). Horizontally, squares are projected onto the continuous decision criterion axis *C*, with *C <* 0 values indicating a tendency toward smile reports and *C >* 0 indicating a tendency toward unsmile reports. Vertically, squares are positioned according to whether the direction of the response-related HFA effect was opposite to, or aligned with, the patient’s decision criterion. In both panels, histograms on the right summarize the number of patients showing opposite, aligned, or tied effects, both bilaterally and separately for the left and right hemispheres.

## Discussion

The present study explored how stimulus-category, response-category, and individual decision criterion effects are reflected in high-frequency activity in human auditory and insular cortices during an ecological voice decision task. Using intracerebral recordings and signal-detection analysis, we found that HFA activity in the STG and insula was more strongly associated with patients’ responses than with the stimulus category.

More importantly, contrary to what would be predicted by a classical hierarchical information-processing view, we found no clear spatial or temporal gradient separating stimulus- and response-related effects; rather, response effects emerged early during stimulus exposure and, strikingly, extended upstream to the most primary sensory regions, namely the transverse gyrus and sulcus of the STG, and posterior regions in the inferior circular sulcus, long gyrus and central sulcus of the insula. Their direction also closely reflected patients’ specific decision criteria.

A central result is the predominance of response-category over stimulus-category effects. Classical feedforward models of information processing (Rauschecker and Tian, 2000; Hickok and Poeppel, 2004; Schirmer and Kotz, 2006) would expect early auditory regions to primarily encode stimulus-related information, with decision-related effects arising later and in higher-order associative regions. Our results showed the opposite tendency: HFA activity in STG and insula was more often associated with the patient’s report than with stimulus category. This held for both overall category contrasts and for SDT-based contrasts in which stimulus category and behavioral report were dissociated. Thus, response-related effects cannot be reduced to acoustic differences between smile and unsmile stimuli, but suggest that these regions tracked perceptual categorization, beyond sensory differences.

These findings are consistent with work showing that sensory cortices are dynamically modulated by task demands, attention and behavioral context (Gilbert and Li, 2013). For instance, primate visual electrophysiology has shown that V1/V2 receptive fields can be modulated by spatial attention (Motter, 1993) or perceptual expectation (Nienborg and Cumming, 2009). However, such effects are usually interpreted as a top-down modulation, with sensory regions acting both as *producers* of information at early latencies and as *receivers* of information later (Cooper et al., 2023). Our results depart from such a hierarchical interpretation in several important aspects.

First, response-related effects were mainly observed during listening, with no evidence that they were preceded by stimulus-related effects in the same or other regions. This rules out the possibility that these response-category effects merely reflected overt reporting or motor preparation, and indicates that activity associated with the eventual report emerged as patients were still processing the auditory sentence, possibly as early as - or *instead of* - activity that encoded stimulus information. Second, although our prefrontal coverage was limited, response-related effects were no *more* prevalent in higher-order regions, such as associative lateral STG, planum temporale or anterior insula, than in primary sensory areas. Together, these results do not suggest a simple, possibly rapid backflow along a cognition-perception axis producing coexisting stimulus and decision representations; rather, decision signals emerge first, and perhaps solely, in areas usually associated with primary sensory processing.

The absence of clear stimulus-category discrimination in auditory regions such as STG and posterior insula is a strong and surprising result. Although this is, to our knowledge, the first iEEG study to investigate smiled-speech perception, the acoustic features distinguishing smiling from non-smiling speech are well described (Tartter, 1980; Ponsot et al., 2018) and overlap with spectral information used to discriminate vowels and other phonetic categories (Strange, 1989). Human intracranial (ECoG) data show that such features are encoded along lateral STG through heterogeneous spatial codes (Oganian et al., 2023), with no clear hemispheric difference and, more generally, that our regions of interest are sensitive to natural speech (Khalighinejad et al., 2021). Here, dominant task-related HFA effects were organized around the patient’s subjective decisions rather than stimulus category, which may result from the ecological nature of the task. Unlike classical paradigms using highly separable stimulus categories, the present task relied on relatively subtle vocal cues embedded in natural speech, making perceptual reports more dependent on internal context and strategy. Methodologically, such ecological tasks may better separate stimulus and decision effects, whereas these may be confounded in highly-separable paradigms. Finally, because much of the literature relies extensively on decoding analyses from neural populations (Napoli et al., 2021; Oganian et al., 2023; Yiling et al., 2024), it may overestimate the decoding separability of stimulus features, especially when responses are heavily biased towards one option, an issue that our single-channel SDT analyses can partly alleviate (see (Soto et al., 2018) for a similar argument).

Several recent intracranial electrophysiology studies have similarly shown that sensory cortices concurrently co-represent stimulus and decision variables in auditory and visual decision tasks (Guo et al., 2019; Napoli et al., 2021; Yiling et al., 2024), sometimes with decision signals in auditory cortex emerging before activity in regions classically associated with decision-making, such as the dorsolateral prefrontal cortex (Napoli et al., 2021). However, almost all such studies were conducted in non-human primates performing highly-constrained learning/reward tasks. To our knowledge, our study is the first to extend such evidence to human participants in an unconditioned, ecological task. A recent human iEEG study attempted to dissociate stimulus and decision features in sensorimotor areas by contrasting report vs. no report tasks using identical tactile stimuli (Albertini et al., 2025), but found that decision-specific activity emerged later and downstream from primary sensorimotor areas – a pattern also found in primate tactile-discrimination studies (Hernandez et al., 2010). This contrast may indicate fundamental differences between tactile and auditory/visual decisions.

More importantly, previous human and monkey studies have remained speculative about why decision- or choice-related information should be back-propagated to sensory areas, and what behavioral relevance these effects may have. Authors have suggested that decision effects in sensory areas may reflect rapid flowback from decision areas, with millisecond conduction delays beyond the temporal resolution of iEEG recordings (Yiling et al., 2024). However, a testable prediction of this view is that decision signals should become available, at similar latencies, across a wide network of interconnected areas, a phenomenon for which we find no convincing evidence here. Alternatively, predictive coding frameworks (Friston, 2005) propose that decision-related signals in sensory areas may encode prediction errors or reward predictions ((Guo et al., 2019); but see Solomon et al. (2021); Westerberg and Roelfsema (2025)). Here again, our paradigm, with randomized stimulus order, balanced distributions and no response feedback, provides no strong support for goal-oriented predictions or rewards.

Instead, in both STG and insula, we found a consistent negative association between response-effect direction and individual decision criteria, in which higher HFA was observed for response categories opposing the patient’s bias. Although descriptive given our small sample, this pattern would be compatible with the idea that sensory HFA reflects the additional sensory evidence required when the report goes against a dominant response tendency. Primary auditory regions would therefore not act as static repositories of stimulus features, but as filters dynamically shaped by task-dependent priors and response strategies. A testable prediction would be that decision effect directionality should dynamically track changes in decision criteria when experimentally manipulated within patients.

Finally, beyond the auditory cortex, our study also provides novel and valuable human iEEG data for the role of posterior and anterior insular cortex in vocal and socio-emotional decision-making. The insula remains relatively poorly understood, but is thought to be involved in detecting salient events and coordinating large-scale behavioral networks (Menon and Uddin, 2010). In auditory contexts, although not generally considered a primary auditory area (but see (Berger et al., 2026)), the insula is thought to obey a posterio-anterior processing axis, with posterior insula showing auditory responses that resemble those in Heschl’s gyrus (Zhang et al., 2019; Berger et al., 2026) whereas anterior areas are associated with decision-making (Loued-Khenissi et al., 2020; Llorens et al., 2023). As in STG, our data largely contradicted this simple hierarchical view: not only did response-related effects in the insula largely predominate, but they also occurred early, flowed back to posterior regions, and were negatively matched to individual decision criteria. Interestingly, these effects also occurred no later than concurrent STG effects, suggesting that the insula participates in forming or stabilizing the perceptual decision in ways that resemble primary auditory cortex, rather than merely reflecting post-decision monitoring or response execution. Taken together, these results prompt further studies of insular contributions to the joint processing of stimulus and decision features, especially in ecological situations where sensory evidence is ambiguous and behaviorally relevant (Rogers-Carter and Christianson, 2019).

In conclusion, the present results provide rare intracerebral evidence that human auditory and insular cortices encode perceptual decisions during an ecological social voice task. Rather than showing a clean separation between early stimulus encoding in sensory cortex and later decision encoding in associative regions, response-related effects emerged early and upstream into the most primary auditory sensory regions. These findings support a view of sensory cortex not as a passive repository of stimulus features, but as an active component of perceptual inference, continuously configured by task demands, priors, and behavioral strategies.

## Author Contributions

CDL designed research, performed research, analyzed data, and wrote the paper; EPR, CF, EL and TS contributed to performing the research by managing or providing access to patients and clinical data; MZ, AM, JP and GD contributed to performing the research by surgically implanting patients; JJA, MG and AL designed research, analyzed data, and wrote the paper. JJA, MG and AL contributed equally as senior authors.

## Acknowledgements

This work was supported by the Agence Nationale de la Recherche (ANR; grant ANR-21-CE19-0048), by the Pôle Neuro Sainte-Anne, GHU Paris Psychiatrie et Neurosciences, through the 2024 Polaire research call funded by intéressement recherche, and by the 2020 “Sauver la Vie” call for projects of the Fondation Université Paris Cité.

