## Supplemental material for "Human primary auditory cortex and insula encode perceptual decisions, not stimulus features"

Table S1: **Patient-level and total anatomical coverage across lobes and major anatomical regions.** Regional columns indicate the number of grey-matter bipolar channels assigned to each region. The Total column indicates the number of unique grey-matter bipolar channels.

| Patient | Temporal lobe | Insular cortex | Frontal lobe | Parietal lobe | Occipital lobe | Total |
|---|---|---|---|---|---|---|
| 1 | 41 | 18 | 63 | 0 | 0 | 118 |
| 2 | 92 | 23 | 4 | 10 | 13 | 136 |
| 3 | 0 | 41 | 43 | 9 | 0 | 87 |
| 4 | 74 | 28 | 32 | 0 | 4 | 130 |
| 5 | 72 | 0 | 0 | 0 | 37 | 105 |
| 6 | 97 | 3 | 11 | 5 | 0 | 114 |
| 7 | 130 | 2 | 0 | 0 | 35 | 162 |
| 8 | 54 | 44 | 19 | 19 | 0 | 131 |
| 9 | 83 | 7 | 0 | 0 | 0 | 87 |
| 10 | 102 | 10 | 0 | 9 | 0 | 117 |
| 11 | 63 | 37 | 62 | 0 | 0 | 158 |
| 12 | 82 | 23 | 26 | 26 | 1 | 152 |
| 13 | 87 | 13 | 20 | 0 | 0 | 115 |
| 14 | 42 | 32 | 52 | 0 | 0 | 122 |
| 15 | 66 | 0 | 4 | 3 | 36 | 99 |
| 16 | 17 | 52 | 29 | 7 | 0 | 93 |
| 17 | 59 | 35 | 38 | 0 | 0 | 123 |
| 18 | 99 | 9 | 0 | 3 | 0 | 106 |
| 19 | 46 | 28 | 31 | 48 | 0 | 149 |
| Total | 1306 | 405 | 434 | 139 | 126 | 2304 |

Table S2: **Behavioral performance and decision criterion by subject.**Behavioral performance and decision criterion by subject. Accuracy corresponds to the percentage of correct responses. Hit, miss, CR, FA percentages are reported relative to the total number of non-neutral trials. Decision criterion $C$ values indicate response tendency: negative values indicate a response tendency toward smile reports, whereas positive values indicate a response tendency toward unsmile reports.

| Patient | Correct (%) | Hit (%) | Miss (%) | CR (%) | FA (%) | $C$ | Response bias |
|---|---|---|---|---|---|---|---|
| 1 | 14.05 | 7.57 | 41.62 | 6.49 | 44.32 | -0.057 | Toward smile |
| 2 | 55.56 | 28.40 | 22.22 | 27.16 | 22.22 | -0.014 | Toward smile |
| 3 | 51.27 | 25.89 | 23.86 | 25.38 | 24.87 | -0.019 | Toward smile |
| 4 | 52.53 | 29.29 | 21.21 | 23.23 | 26.26 | -0.138 | Toward smile |
| 5 | 66.16 | 30.81 | 19.70 | 35.35 | 14.14 | 0.142 | Toward unsmile |
| 6 | 56.50 | 14.12 | 31.64 | 42.37 | 11.86 | 0.630 | Toward unsmile |
| 7 | 51.81 | 23.32 | 26.42 | 28.50 | 21.76 | 0.122 | Toward unsmile |
| 8 | 51.26 | 10.55 | 39.70 | 40.70 | 9.05 | 0.846 | Toward unsmile |
| 9 | 56.98 | 39.53 | 12.79 | 17.44 | 30.23 | -0.511 | Toward smile |
| 10 | 58.64 | 30.89 | 18.85 | 27.75 | 22.51 | -0.088 | Toward smile |
| 11 | 66.32 | 32.64 | 17.62 | 33.68 | 16.06 | 0.037 | Toward unsmile |
| 12 | 49.24 | 24.87 | 24.87 | 24.37 | 25.89 | -0.019 | Toward smile |
| 13 | 52.29 | 25.49 | 22.88 | 26.80 | 24.84 | -0.010 | Toward smile |
| 14 | 61.02 | 3.39 | 37.29 | 57.63 | 1.69 | 1.507 | Toward unsmile |
| 15 | 68.75 | 39.38 | 8.12 | 29.38 | 23.12 | -0.393 | Toward smile |
| 16 | 65.66 | 39.90 | 9.60 | 25.76 | 24.75 | -0.414 | Toward smile |
| 17 | 70.47 | 35.75 | 13.99 | 34.72 | 15.54 | -0.040 | Toward smile |
| 18 | 50.50 | 27.00 | 23.00 | 23.50 | 26.50 | -0.087 | Toward smile |
| 19 | 58.08 | 38.38 | 11.62 | 19.70 | 30.30 | -0.494 | Toward smile |

Table S3: **Subject-level distribution of task-responsive channels in the STG and insula by hemisphere and task period.** Values are reported as percentages with raw counts in parentheses, where the denominator is the number of tested channels for each subject, hemisphere, and region. A dash indicates that no channel was tested for that subject, hemisphere, and region.

| Subject | Hemisphere | ROI | Tested channels | TR any | TR listening | TR delay | TR response |
|---|---|---|---|---|---|---|---|
| 1 | Left | STG | 0 | – | – | – | – |
| | | Insula | 0 | – | – | – | – |
| | Right | STG | 4 | 75.0% (3/4) | 50.0% (2/4) | 50.0% (2/4) | 25.0% (1/4) |
| | | Insula | 18 | 55.6% (10/18) | 50.0% (9/18) | 11.1% (2/18) | 5.6% (1/18) |
| 2 | Left | STG | 0 | – | – | – | – |
| | | Insula | 0 | – | – | – | – |
| | Right | STG | 38 | 73.7% (28/38) | 73.7% (28/38) | 39.5% (15/38) | 0.0% (0/38) |
| | | Insula | 23 | 34.8% (8/23) | 30.4% (7/23) | 4.3% (1/23) | 4.3% (1/23) |
| 3 | Left | STG | 0 | – | – | – | – |
| | | Insula | 0 | – | – | – | – |
| | Right | STG | 0 | – | – | – | – |
| | | Insula | 41 | 19.5% (8/41) | 12.2% (5/41) | 2.4% (1/41) | 7.3% (3/41) |
| 4 | Left | STG | 29 | 69.0% (20/29) | 69.0% (20/29) | 62.1% (18/29) | 3.4% (1/29) |
| | | Insula | 28 | 42.9% (12/28) | 21.4% (6/28) | 28.6% (8/28) | 32.1% (9/28) |
| | Right | STG | 0 | – | – | – | – |
| | | Insula | 0 | – | – | – | – |
| 5 | Left | STG | 0 | – | – | – | – |
| | | Insula | 0 | – | – | – | – |
| | Right | STG | 10 | 10.0% (1/10) | 10.0% (1/10) | 0.0% (0/10) | 0.0% (0/10) |
| | | Insula | 0 | – | – | – | – |
| 6 | Left | STG | 9 | 77.8% (7/9) | 77.8% (7/9) | 77.8% (7/9) | 0.0% (0/9) |
| | | Insula | 2 | 100.0% (2/2) | 100.0% (2/2) | 100.0% (2/2) | 0.0% (0/2) |
| | Right | STG | 19 | 57.9% (11/19) | 57.9% (11/19) | 47.4% (9/19) | 0.0% (0/19) |
| | | Insula | 1 | 100.0% (1/1) | 100.0% (1/1) | 100.0% (1/1) | 0.0% (0/1) |
| 7 | Left | STG | 17 | 41.2% (7/17) | 35.3% (6/17) | 17.6% (3/17) | 29.4% (5/17) |
| | | Insula | 0 | – | – | – | – |
| | Right | STG | 24 | 54.2% (13/24) | 37.5% (9/24) | 0.0% (0/24) | 25.0% (6/24) |
| | | Insula | 2 | 0.0% (0/2) | 0.0% (0/2) | 0.0% (0/2) | 0.0% (0/2) |
| 8 | Left | STG | 41 | 85.4% (35/41) | 85.4% (35/41) | 29.3% (12/41) | 14.6% (6/41) |
| | | Insula | 44 | 43.2% (19/44) | 36.4% (16/44) | 13.6% (6/44) | 6.8% (3/44) |
| | Right | STG | 0 | – | – | – | – |
| | | Insula | 0 | – | – | – | – |
| 9 | Left | STG | 23 | 26.1% (6/23) | 21.7% (5/23) | 21.7% (5/23) | 4.3% (1/23) |
| | | Insula | 7 | 42.9% (3/7) | 42.9% (3/7) | 28.6% (2/7) | 0.0% (0/7) |
| | Right | STG | 0 | – | – | – | – |
| | | Insula | 0 | – | – | – | – |
| 10 | Left | STG | 26 | 53.8% (14/26) | 46.2% (12/26) | 50.0% (13/26) | 15.4% (4/26) |
| | | Insula | 10 | 40.0% (4/10) | 30.0% (3/10) | 30.0% (3/10) | 0.0% (0/10) |
| | Right | STG | 0 | – | – | – | – |
| | | Insula | 0 | – | – | – | – |
| 11 | Left | STG | 18 | 38.9% (7/18) | 27.8% (5/18) | 33.3% (6/18) | 5.6% (1/18) |
| | | Insula | 23 | 60.9% (14/23) | 56.5% (13/23) | 47.8% (11/23) | 4.3% (1/23) |
| | Right | STG | 2 | 0.0% (0/2) | 0.0% (0/2) | 0.0% (0/2) | 0.0% (0/2) |
| | | Insula | 14 | 71.4% (10/14) | 57.1% (8/14) | 57.1% (8/14) | 14.3% (2/14) |
| 12 | Left | STG | 37 | 67.6% (25/37) | 64.9% (24/37) | 56.8% (21/37) | 5.4% (2/37) |

*Continued on next page*

| Subject | Hemisphere | ROI | Tested channels | TR any | TR listening | TR delay | TR response |
|---|---|---|---|---|---|---|---|
| | | Insula | 23 | 17.4% (4/23) | 17.4% (4/23) | 13.0% (3/23) | 4.3% (1/23) |
| | Right | STG | 0 | – | – | – | – |
| | | Insula | 0 | – | – | – | – |
| 13 | Left | STG | 35 | 54.3% (19/35) | 51.4% (18/35) | 40.0% (14/35) | 2.9% (1/35) |
| | | Insula | 13 | 23.1% (3/13) | 23.1% (3/13) | 23.1% (3/13) | 0.0% (0/13) |
| | Right | STG | 0 | – | – | – | – |
| | | Insula | 0 | – | – | – | – |
| 14 | Left | STG | 0 | – | – | – | – |
| | | Insula | 0 | – | – | – | – |
| | Right | STG | 17 | 88.2% (15/17) | 88.2% (15/17) | 64.7% (11/17) | 0.0% (0/17) |
| | | Insula | 32 | 18.8% (6/32) | 15.6% (5/32) | 3.1% (1/32) | 3.1% (1/32) |
| 15 | Left | STG | 1 | 100.0% (1/1) | 100.0% (1/1) | 0.0% (0/1) | 0.0% (0/1) |
| | | Insula | 0 | – | – | – | – |
| | Right | STG | 20 | 80.0% (16/20) | 80.0% (16/20) | 30.0% (6/20) | 15.0% (3/20) |
| | | Insula | 0 | – | – | – | – |
| 16 | Left | STG | 17 | 88.2% (15/17) | 88.2% (15/17) | 70.6% (12/17) | 0.0% (0/17) |
| | | Insula | 52 | 48.1% (25/52) | 36.5% (19/52) | 23.1% (12/52) | 19.2% (10/52) |
| | Right | STG | 0 | – | – | – | – |
| | | Insula | 0 | – | – | – | – |
| 17 | Left | STG | 39 | 35.9% (14/39) | 33.3% (13/39) | 28.2% (11/39) | 2.6% (1/39) |
| | | Insula | 27 | 40.7% (11/27) | 22.2% (6/27) | 22.2% (6/27) | 22.2% (6/27) |
| | Right | STG | 0 | – | – | – | – |
| | | Insula | 8 | 37.5% (3/8) | 0.0% (0/8) | 0.0% (0/8) | 37.5% (3/8) |
| 18 | Left | STG | 33 | 69.7% (23/33) | 66.7% (22/33) | 51.5% (17/33) | 6.1% (2/33) |
| | | Insula | 9 | 44.4% (4/9) | 44.4% (4/9) | 33.3% (3/9) | 11.1% (1/9) |
| | Right | STG | 0 | – | – | – | – |
| | | Insula | 0 | – | – | – | – |
| 19 | Left | STG | 0 | – | – | – | – |
| | | Insula | 0 | – | – | – | – |
| | Right | STG | 28 | 57.1% (16/28) | 53.6% (15/28) | 32.1% (9/28) | 3.6% (1/28) |
| | | Insula | 28 | 39.3% (11/28) | 32.1% (9/28) | 10.7% (3/28) | 3.6% (1/28) |
